# Neutralizing antibody 5-7 defines a distinct site of vulnerability in SARS-CoV-2 spike N-terminal domain

**DOI:** 10.1101/2021.06.29.450397

**Authors:** Gabriele Cerutti, Yicheng Guo, Pengfei Wang, Manoj S. Nair, Yaoxing Huang, Jian Yu, Lihong Liu, Phinikoula S. Katsamba, Fabiana Bahna, Eswar R. Reddem, Peter D. Kwong, David D. Ho, Zizhang Sheng, Lawrence Shapiro

**Author notes:** Equal contribution (Gabriele Cerutti, Yicheng Guo). Correspondence (D.D.H.), (Z.S.), (L.S.).

## Abstract

Antibodies that potently neutralize SARS-CoV-2 target mainly the receptor-binding domain or the N-terminal domain (NTD). Over a dozen potently neutralizing NTD-directed antibodies have been studied structurally, and all target a single antigenic supersite in NTD (site 1). Here we report the 3.7 Å resolution cryo-EM structure of a potent NTD-directed neutralizing antibody 5-7, which recognizes a site distinct from other potently neutralizing antibodies, inserting a binding loop into an exposed hydrophobic pocket between the two sheets of the NTD β-sandwich. Interestingly, this pocket has been previously identified as the binding site for hydrophobic molecules including heme metabolites, but we observe their presence to not substantially impede 5-7 recognition. Mirroring its distinctive binding, antibody 5-7 retains a distinctive neutralization potency with variants of concern (VOC). Overall, we reveal a hydrophobic pocket in NTD proposed for immune evasion can actually be used by the immune system for recognition.

**Highlights:**

- Cryo-EM structure of neutralizing antibody 5-7 in complex with SARS CoV-2 spike
- 5-7 recognizes NTD outside of the previously identified antigenic supersite
- 5-7 binds to a site known to accommodate numerous hydrophobic ligands
- Structural basis of 5-7 neutralization tolerance to some variants of concern

## Introduction

Severe acute respiratory syndrome coronavirus 2 (SARS-CoV-2), the causative agent for Coronavirus Disease 2019 (COVID-19), emerged in 2019, leading to the ongoing worldwide pandemic which has now led to over three million deaths (Callaway et al., 2020; Cucinotta and Vanelli, 2020; Dong et al., 2020). Over 300 million COVID-19 cases have been reported, giving ample opportunity for the emergence of mutants, and numerous variants of concern (VOC) including those prevalent the UK (B.1.1.7), South Africa (B.1.351), Brazil (P. 1), New York City (B.1.526), California (B.1.427/9), and India (B.1.617) have become dominant in some geographic regions and are now driving the pandemic (Annavajhala et al., 2021; Nuno R. Faria et al., 2021; Tang et al., 2021; Tegally et al., 2020; Yadav et al., 2021; Zhang et al., 2021). Numerous mutations in circulating VOC are within the epitopes of neutralizing antibody classes common in the convalescent human repertoire, consistent with their selection in response to human immune pressure (Cerutti et al., 2021b; Rapp et al., 2021; Yuan et al., 2020). While effective vaccines are now being widely distributed in some countries, the pace and geographic restriction of vaccine administration ensures the continued rise of new infections and the continued need for therapeutics.

One promising therapeutic approach – the identification of SARS-CoV-2-neutralizing antibodies that could be used as therapeutic or prophylactic agents – has now been extensively explored. The primary target for neutralizing antibodies is the viral spike protein, a trimeric type I viral fusion machine (Walls et al., 2020; Wrapp et al., 2020b) which allows the virus to bind to the ACE2 receptor on host cells through its receptor-binding domain (RBD) (Benton et al., 2020; Yan et al., 2020; Zhou et al., 2020) and mediates fusion between the viral and cell membranes. The spike protein is comprised of two subunits: the S1 subunit comprising the N-terminal domain (NTD), RBD and several other subdomains, and the S2 subunit that mediates virus–cell membrane fusion (Walls et al., 2020; Wrapp et al., 2020b). The majority of SARS-CoV-2 neutralizing antibodies so far identified target RBD (Brouwer et al., 2020; Cao et al., 2020; Chen et al., 2020; Chi et al., 2020; Ju et al., 2020; Liu et al., 2020b; Pinto et al., 2020; Robbiani et al., 2020; Rogers et al., 2020; Seydoux et al., 2020; Wang et al., 2020a; Wrapp et al., 2020a; Wu et al., 2020b; Zeng et al., 2020; Zost et al., 2020). Structural studies (Barnes et al., 2020a; Barnes et al., 2020b; Liu et al., 2020a; Wang et al., 2020b; Yuan et al., 2020) have revealed neutralizing antibodies to recognize RBD at multiple distinct sites, and further revealed multi-donor RBD-directed antibody classes that appear to be elicited with high frequency in the human population (Barnes et al., 2020b; Robbiani et al., 2020; Wu et al., 2020a; Yuan et al., 2020). Neutralization for many RBD-directed antibodies can be explained by interference with RBD-ACE2 interaction, and/or impeding the ability of RBD to adopt the “up” conformation (Barnes et al., 2020b; Liu et al., 2020a; Yuan et al., 2020) required for ACE2 binding (Benton et al., 2020).

The most effective NTD-directed neutralizing antibodies have potencies rivaling those of the best RBD-directed neutralizing antibodies, and recent studies have extensively characterized the NTD-directed antibody response for SARS-CoV-2. Veesler and co-workers isolated a panel of 41 NTD-directed antibodies and mapped their recognition to three sites (McCallum et al., 2021b). Remarkably, all antibodies with neutralizing activity mapped to a single site, site 1. In a parallel study, we determined structures for seven NTD-directed antibodies selected for their potent neutralizing activities, finding them all to target a single supersite that coincides with site 1 identified by Veesler (Cerutti et al., 2021a). Both papers designated the site 1 region as an “antigenic supersite”, targeted by diverse lineages of neutralizing antibodies. In addition to these studies, Crowe and coworkers determined structures of two NTD-directed neutralizing antibodies, both of which targeted the site 1 supersite (Suryadevara et al., 2021). Four additional cryo-EM structures for SARS-CoV-2 NTD-directed neutralizing antibodies - 4A8, FC05, CM25, and DH1050.1, reveal binding to the site 1 antigenic supersite (Chi et al., 2020; Li et al., 2021; Voss et al., 2021; Wang et al., 2021a), while a single structure for a non-neutralizing antibody, DH1052, shows targeting outside the supersite (Li et al., 2021). Overall, these results supported the idea that the site 1 supersite represented the lone site of neutralization vulnerability in NTD, which results in high selection pressure for viral escape.

Structural studies have characterized the NTD antigenic supersite and other nonneutralizing epitopes on NTD (**Figure S1**). NTD is highly glycosylated, and the site 1 supersite, located at the periphery of the spike, is the largest glycan-free surface on NTD facing away from the viral membrane (Cerutti et al., 2021a). Antibodies bind to a region of flexible loops, centered on the N3 β-hairpin. The supersite is highly electropositive. Epitopes to other less potent or non-neutralizing antibodies have also been mapped and numbered as sites 2 through 6. The basis for the restriction of potent neutralization to site 1 is unclear, however preliminary studies are consistent with a conformational mechanism for supersite-directed antibodies in which membrane fusion is inhibited (McCallum et al., 2021b; Suryadevara et al., 2021). Despite the potent neutralization of the initial 2019 strain of SARS-CoV-2 achieved by neutralizing antibodies, numerous VOC have emerged, some of which show concerning resistance to neutralizing antibodies. Antibodies that target the NTD antigenic supersite appear to be particularly vulnerable to escape, with many losing all activity against the UK, South Africa, and Brazil VOC (Wang et al., 2021b; Wang et al., 2021c).

Here we describe the cryo-EM structure of antibody 5-7 in complex with SARS-CoV-2 spike, present neutralization data with VOC, and analyze the recognized epitope. We show that the 5-7 binding site represents a second site of neutralization vulnerability in SARS CoV-2 NTD remote from most VOC mutations, underscoring the potential therapeutic value of antibody 5-7.

## Results

### Antibody 5-7 targets a hydrophobic pocket in NTD

From a complex of SARS-CoV-2 spike - stabilized by 2P mutations (Wrapp et al., 2020b) - with the antigen-binding fragment (Fab) of antibody 5-7, we collected single particle data on a Titan Krios microscope, yielding a cryo-EM reconstruction to an overall resolution of 3.70 Å (**Figure 1A, Figures S1 and S2, Table S1**). The major 3D class had one Fab per trimer. Recognition of NTD by 5-7 was dominated by the heavy chain, which buried 1169.3 Å^2^ surface area, with a smaller 163.3 Å^2^ contribution by the light chain. The 24 amino acid CDR H3 formed the dominant interaction with NTD, with additional contributions from CDRs H2, L1 and L3 (**Figure 1B; Figure S3**). CDR H3 formed hydrophobic and hydrogen bond contacts with NTD, in particular through residues near the tip of the CDR H3 loop, W1000f, S100h, and L100i, which are within a short helix. Further toward the base of CDR H3, four hydrogen bonds are formed with NTD and additional hydrophobic contacts are formed, particularly with the side chain of CDR H3 Y100e. CDR H2 contributed to NTD recognition mainly through hydrogen bonds formed between the hydroxyl groups of Ser53 and Ser56. Light chain interacted with NTD residues Q173 and P174 through hydrogen bond and hydrophobic interactions, respectively (**Figure S3A**).

**Figure 1.**
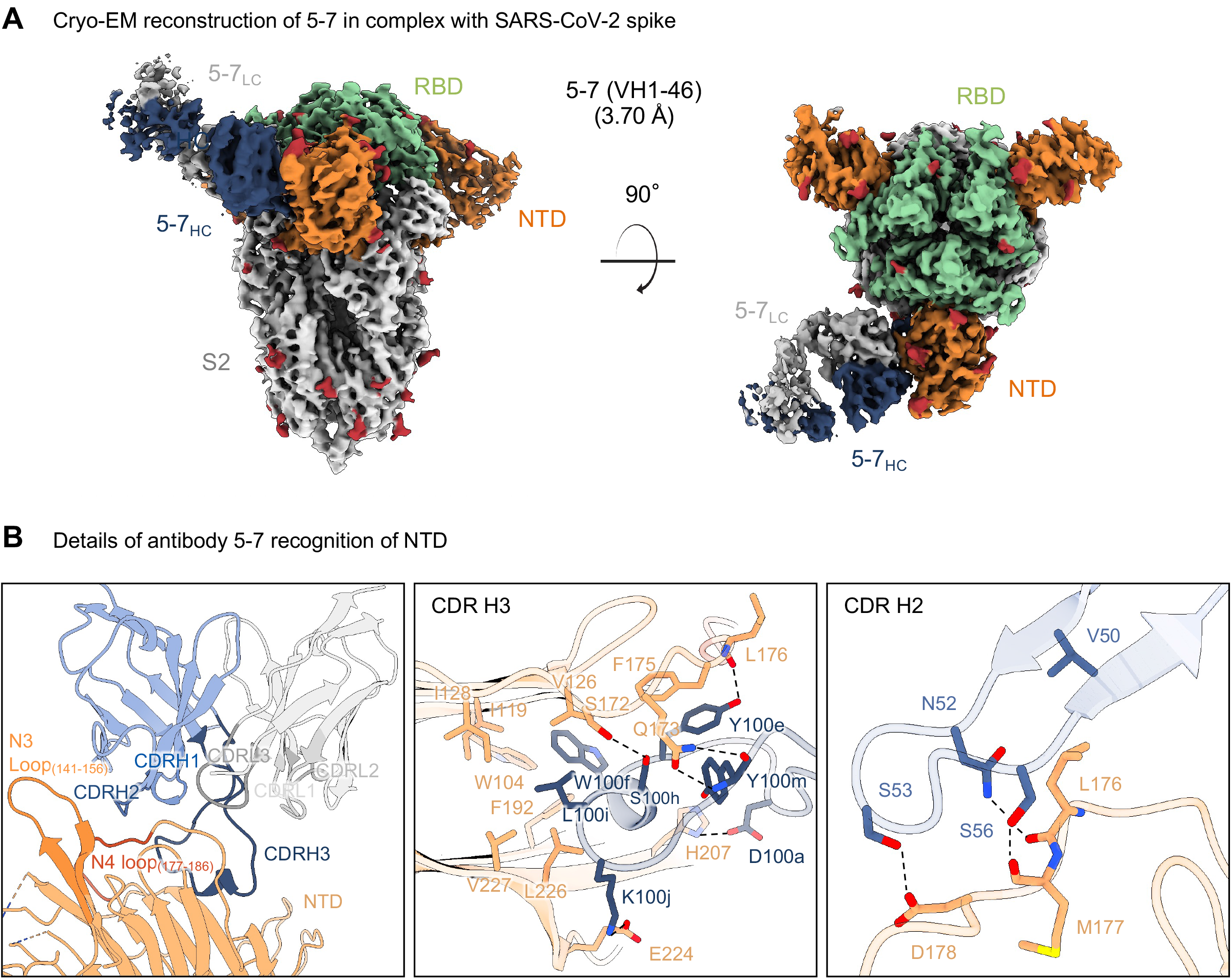
Antibody 5-7 targets a hydrophobic pocket in NTD. (A) Cryo-EM reconstruction for spike complex with antibody 5-7 from two orthogonal views. NTD is shown in orange, RBD in green, glycans in red, antibody heavy chain in blue and light chain in gray. (B) Details of antibody 5-7 recognition of NTD showing the overall interface (left panel), recognition by CDR H3 (middle panel) and recognition by CDR H2 (right panel). CDR H1, H2, H3 are colored in shades of blue; CDR L1, L2, and L3 are colored in shades of gray. Nitrogen atoms are colored in blue, oxygen atoms in red; hydrogen bonds (distance <3.2 Å) are represented as dashed lines. See also Figures S1, S2 and S3 and Table S1.

### Antibody 5-7 targets a site distinct from the NTD supersite

Since other neutralizing antibodies targeting NTD recognize the site 1 antigenic supersite, we compared the recognition of NTD by 5-7 and supersite antibodies. We produced a structural superposition of all NTD-directed antibodies deposited in the PDB, superposed on NTD Ca atoms, in the context of SARS-CoV-2 spike trimer (**Figure 2A**). All neutralizing antibodies target the NTD supersite except for 5-7. The single non-neutralizing antibody DH1052 binds to a different region at the “bottom” of NTD.

**Figure 2.**
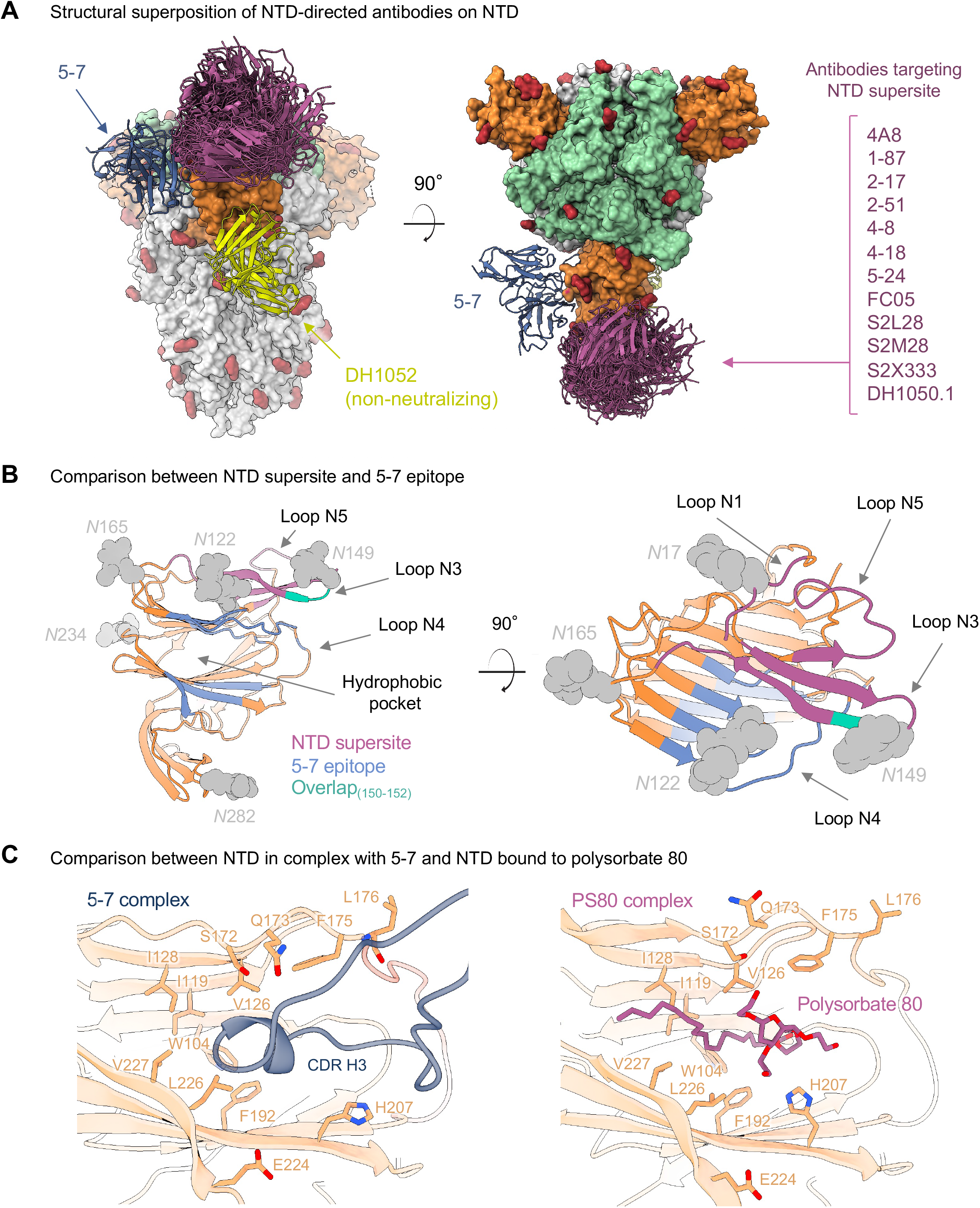
Antibody 5-7 adopts a different binding mode compared to antibodies targeting the NTD supersite. (A) Structural superposition of all NTD-directed antibodies deposited in the PDB on NTD in the context of SARS-CoV-2 spike trimer. All neutralizing antibodies target the NTD supersite (magenta) except for 5-7 (blue). The non-neutralizing antibody DH1052 (yellow) binds to a different region at the “bottom” of NTD. (B) Comparison between NTD supersite (magenta) and 5-7 epitope (blue) on NTD. The overlap region only includes three residues (150-152) located in the N3 loop. (C) The hydrophobic pocket that accommodates 5-7 CDR H3 (left panel) has been previously reported to bind polysorbate 80 (right panel). See also Figure S3.

Overall, the 5-7 recognition site and NTD supersite (as defined by Cerutti et al., 2021a) are distinct, with a small area of overlap. This can be seen in the ribbon diagram in **Figure 2B** comparing the NTD supersite shown in magenta with the 5-7 epitope shown in blue. The overlap region only includes three residues (150-152) located in the N3 loop. Since the supersite was defined for multiple antibodies of different classes, it is larger than the footprint of any single supersite antibody. Nevertheless, of the 12 supersite antibodies we assessed, all of them except S2L28 show direct van der Waals clashes with 5-7, localized either to the N3 β-harpin or in the N4 loop region.

The hydrophobic pocket that accommodates 5-7 CDR H3 has been previously reported to bind hydrophobic ligands, including polysorbate 80 (Bangaru et al., 2020) and biliverdin (Rosa et al., 2021). Comparison of the structure of NTD in complex with polysorbate 80 with the structure of the 5-7 NTD complex revealed nearly identical recognition within the hydrophobic pocket (**Figure 2C**). It has been suggested that such ligands could prevent binding of supersite antibodies by conformational competition (Rosa et al., 2021). We therefore assessed binding of 5-7 and supersite antibodies 4-8 and 5-24 to NTD in the presence of these small molecules (**Figure S4**). We ran SPR binding experiments in the presence of biliverdin, bilirubin and polysorbate 80 – all reported to bind to the 5-7 pocket – with each at a concentration greater than natural abundance (Shum et al., 2021) and greater than 10-fold its K_D_ for binding NTD. In each case binding was observed, despite attenuated affinity in the case of biliverdin for 5-7 binding NTD.

### Neutralization by antibody 5-7 is partially tolerant to B.1.1.7, B.1.351, and B.1.526 variants

We assessed neutralization of five variants of SARS-CoV-2 infectious virus using three antibodies directed against NTD domain (**Figure 3A**). While the potency of all the NTD monoclonal antibodies (mAbs) were lowered against the variants in comparison to the original isolate (WA1), we found that only one of the mAbs, 5-7, retained ~50% of the neutralization activity to all the examined variants. This difference in functional behavior of 5-7 against the live viruses was dissected by generating the key NTD mutations into VSV-based pseudovirus followed by testing the neutralization of each virus. 5-7 showed very different neutralization profiles as compared to the two NTD supersite targeting antibodies, 5-24 and 4-8. While 5-24 and 4-8 completely lost their activities against the mutations fall in the NTD supersite (144del, W152C, 242-244del and R246I), 5-7 still retained its activity, at least partially, against those mutations (**Figure 3B**).

**Figure 3.**
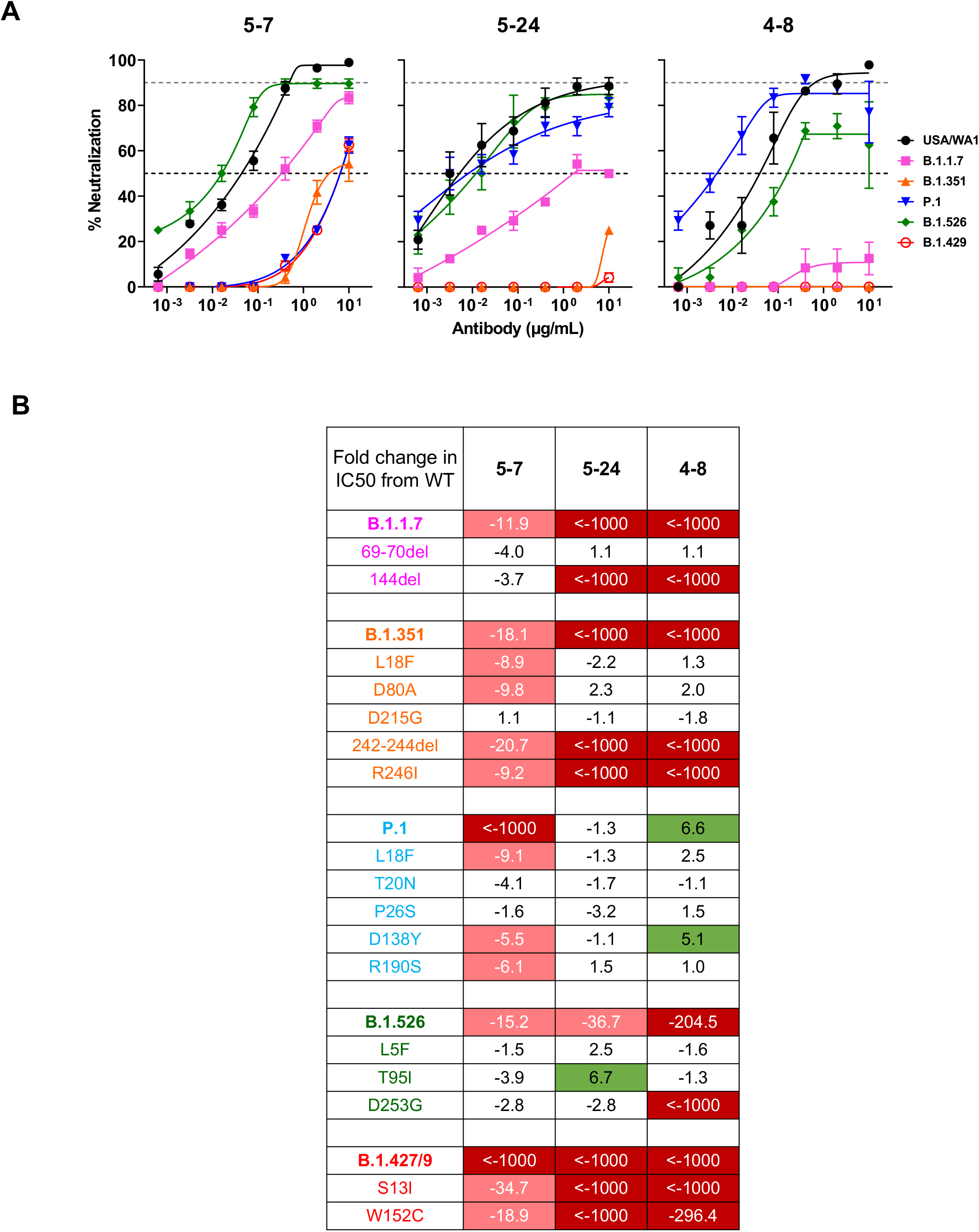
SARS-CoV-2 neutralization profiles for 5-7 and two NTD supersite targeting antibodies. (A) Neutralization of wild-type (USA/WA1), B.1.1.7, B.1.351, P.1, B.1.526 and B.1.429 authentic viruses by NTD-directed mAbs 5-7, 5-24 and 4-8. The horizontal dotted lines on each graph indicate 50% and 90% neutralization. Data are mean ± s.e.m. of technical triplicates and represent one of two independent experiments. (B) Fold increase or decrease in IC50 of the mAbs against pseudoviruses containing all the combined mutations or the NTD single-mutation of B.1.1.7, B.1.351, P.1, B.1.526 and B.1.427/9, relative to the wild-type (D614G) virus, presented as a heat map in which darker colors indicate a greater change. Red, resistance >5-fold; green, sensitization >5-fold.

### The epitope of antibody 5-7 is remote from most VOC mutations

While potency was reduced, antibody 5-7 retained ~50% neutralization activity to all the variants examined as live virus. To understand the basis for this tolerance, we mapped the mutations of each variant – UK (B.1.1.7), South Africa (B.1.351), Brazil (P.1), New York (B.1.561), and California (B.1.427/9) – to the surface of NTD in the spike complex structure with 5-7 we report here (**Figure 4A**). Of the 17 mutations, L5F and S13I located within the signal peptide, 8 mapped within the footprint of the supersite. The epitope of antibody 5-7 contains 2 VOC mutants, R190S in the Brazil P.1 virus and W152C in the California B.1.427/9 variant. The structure modeling reveals that both R190S and W152C impair 5-7 binding by altering the local conformation of NTD loops (**Figure S3B**). R190S and W152C each attenuates pseudovirus IC50 by 6.1-fold and 18.9-fold, respectively. We also observed NTD mutations far from the 5-7 epitope attenuate 5-7 potency (**Figure 3B**), suggesting that other mutations may remotely modulate the conformation of NTD, a mechanism similar to other NTD-directed antibodies (McCallum et al., 2021a; Wang et al., 2021b). This agrees with the more substantial attenuation of 5-7 potency observed with live virus neutralization for the P.1 and B.1.427/9 strains (**Figure 3A**).

**Figure 4.**
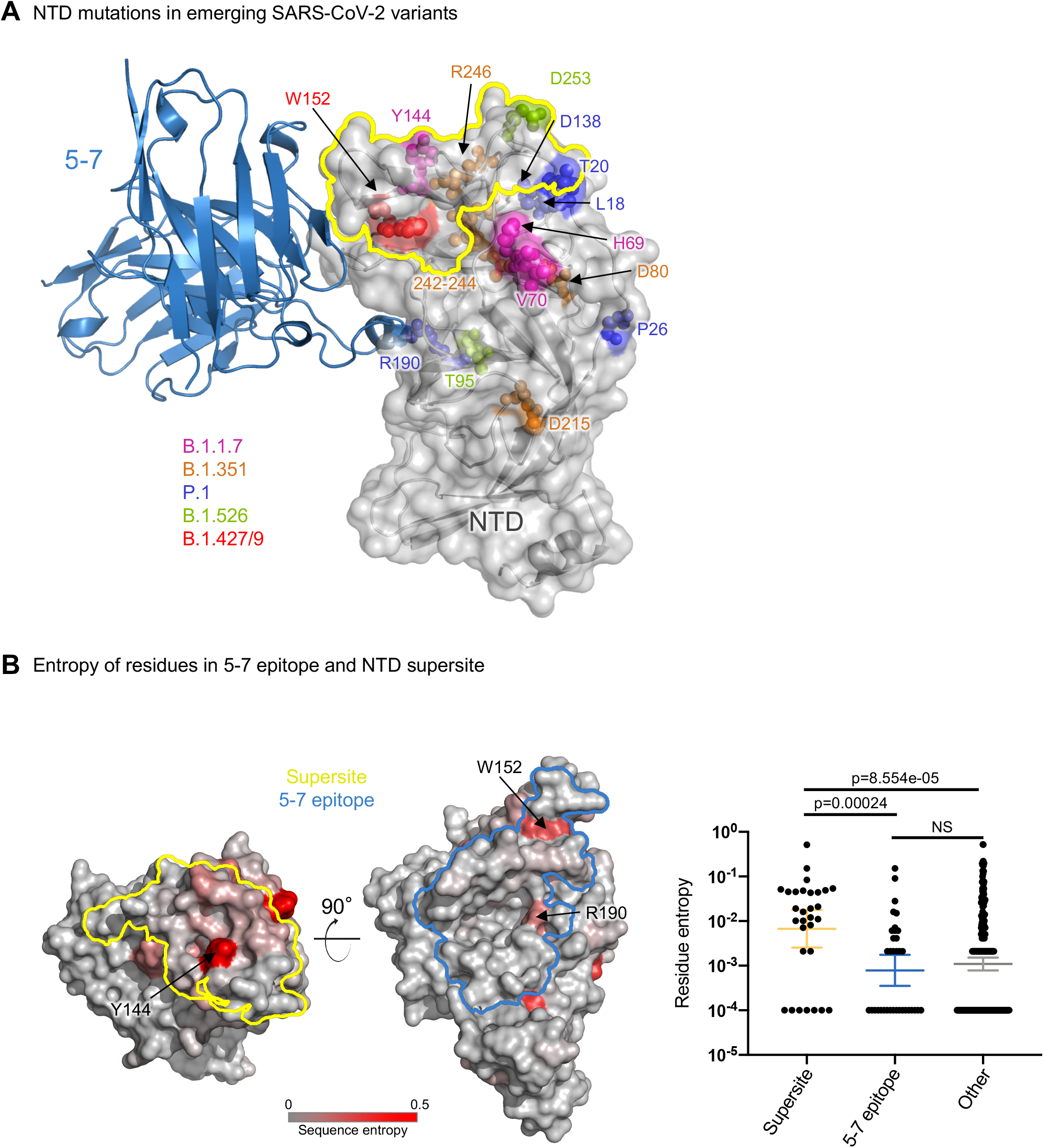
Antibody 5-7 binds apart from most of the mutations in emerging SARS-CoV-2 variants. (A) The majority of NTD mutations in different SARS-CoV-2 variants are not within the 5-7 binding region. Mutations in B.1.1.7 are colored magenta, B.1.351 orange, P.1 blue, B.1.526 green, and B.1.427/9 red respectively. The NTD supersite is outlined in yellow. (B) Comparison of residue entropy between NTD supersite, 5-7 epitope and other residues on NTD. Larger entropy means more diversification. Left and middle panels show the footprint and residue entropy for supersite and 5-7 epitope on NTD surface. Right panel: residues are represented as dots, the geometric mean and 95% CI of the three groups of residues were showed in blue, red and green, respectively. 5-7 epitope residues are the most conserved among the three groups. The p values between different groups were calculated using the Kolmogorov–Smirnov test. See also Figure S3.

To assess the potential breadth of 5-7, we compared the sequence entropy of its epitope among SARS-CoV-2 sequences in the GISAID database to the sequence entropy of the supersite targeted by neutralizing antibodies, and the rest of the NTD surface (**Figure 4B**). The sequence entropy of the 5-7 epitope is about an order of magnitude lower than that of the supersite. Not surprisingly, the W152 and R190 showed the highest entropy within the 5-7 interface, the other residues are highly conserved. Overall, this shows that the 5-7 epitope has low variability, and suggests that 5-7 can successfully target numerous circulating SARS-CoV-2 variants.

## Discussion

The structure of 5-7 in complex with SARS-CoV-2 spike shows that potent neutralization can be achieved by antibodies that target NTD outside of the site 1 supersite. In vitro experiments suggest that supersite-directed antibodies neutralize not by blocking recognition of the ACE2 receptor, but by inhibiting conformational changes required for fusion (Suryadevara et al., 2021). It is possible that 5-7, which binds at a site near to but distinct from the supersite and also fails to inhibit interaction with ACE2 (Liu et al., 2020a), could function through a similar mechanism.

Despite the clear structural uniqueness of antibody 5-7, prior experiments showed binding competition between 5-7 and supersite-directed antibodies (Liu et al., 2020a). The strong competition observed between 5-7 and all supersite antibodies could be steric, but could also involve conformational competition: comparison of the NTD conformations observed in each antibody revealed a structural coupling between the N3 β-harpin and the N4 loop outlining the epitope of 5-7, which acts as a gate for the hydrophobic pocket (**Figure S3C**). In most antibody-bound structures the N4 loop adopts a “closed” conformation, preventing exposure of the hydrophobic pocket, but the “open” conformation is required to accommodate the CDR H3 loop of 5-7. The N4 loop in the S2L28-bound NTD structure was modelled in the “open” conformation, thus resembling the conformation observed in the structure of 5-7-bound NTD, but the EM density shows an equilibrium of both the “open” and “closed” states. As a result, the NTD conformation observed in the 5-7 complex structure is unique and appears to be incompatible with binding of other reported NTD-directed neutralizing antibodies.

The site that accommodates CDR H3, between the two sheets of the NTD β-sandwich, has also been shown to accommodate small molecule ligands, including a detergent, polysorbate 80 (Bangaru et al., 2020), and heme metabolites biliverdin and bilirubin (Rosa et al., 2021). The SPR binding experiments reported here show that binding of these compounds does not significantly interfere with recognition by 5-7 or supersite antibodies 5-24 and 4-8 (**Figure S4**). This hydrophobic pocket may play a role in SARS-CoV-2 biology, such as interacting with a target protein ligand, but if so, this remains unknown. The binding of biliverdin and bilirubin has been proposed as a means of viral evasion (Rosa et al., 2021), though we find here the immune system is able to generate an antibody that turns this binding pocket into a site of vulnerability for antibody neutralization.

Among the more attractive properties of antibody 5-7 is the partial tolerance of its neutralization to VOC mutations. We have shown here that this tolerance arises largely due to its novel binding orientation, which leads to a lack of overlap of 5-7 with VOC mutations. Overall, we have shown 5-7 to target a conserved epitope on NTD, increasing the number of known neutralizing epitopes and enabling therapeutic strategies such as cocktail combinations or multispecifics that target both NTD and RBD.

## Acknowledgments

We thank R. Grassucci, Y.-C. Chi and Z. Zhang from the Cryo-EM Center at Columbia University for assistance with cryo-EM data collection. Support for this work was also provided by the Intramural Program of the Vaccine Research Center, National Institutes of Allergy and Infectious Diseases, NIH, and by the Samuel Yin, Pony Ma, Peggy & Andrew Cherng, Brii Bioscieces, Jack Ma Foundation, JBP Foundation, Carol Ludwig, and Roger & David Wu, COVID-19 Fast Grants, the Self Graduate Fellowship Program, and NIH grants DP5OD023118, R21AI143407, and R21AI144408.

## Author Contributions

GC determined the cryo-EM structure of 5-7, YG performed bioinformatics analysis. PW, MSN, YH, JY, LL, DDH contributed data on neutralization assays and also produced IgGs and Fabs. FB produced spike. JY and ERR produced Fabs. DDH supervised IgG production and neutralization assay, PDK contributed to bioinformatics and structural analysis, ZS supervised bioinformatics analysis, LS supervised structural determinations and led the overall project. GC, YG, PDK, ZS and LS wrote the manuscript, with all authors comments.

## STAR★METHODS

- KEY RESOURCES TABLE
- RESOURCE AVAILABILITY

- Lead contact
- Materials availability
- Data and code availability
- EXPERIMENTAL MODEL AND SUBJECT DETAILS

- Cell lines
- METHOD DETAILS

- Protein samples expression and purification
- Cryo-EM sample preparation
- Cryo-EM data collection, processing and structure refinement
- Authentic SARS-CoV-2 microplate neutralization
- Pseudovirus neutralization assays
- SPR experiments
- Antibody gene assignments and genetic analyses
- Sequence entropy comparison
- QUANTIFICATION AND STATISTICAL ANALYSIS

### RESOURCE AVAILABILITY

#### Lead contact

Further information and requests for resources and reagents should be directed to and will be fulfilled by Lawrence Shapiro.

#### Materials availability

This study did not generate new unique reagents.

#### Data and code availability

The cryo-EM structures have been deposited to the Electron Microscopy Data Bank (EMDB) and the Protein Data Bank (RCSB PDB). Cryo-EM structural models and maps for antibody 5-7 in complex with SARS-CoV-2 spike have been deposited in the PDB and EMDB with accession codes PDB 7N01, EMD-24097 respectively.

### EXPERIMENTAL MODEL AND SUBJECT DETAILS

#### Cell lines

FreeStyle 293-F (cat# R79007), Expi293F cells (cat# A14635) were from Thermo Fisher Scientific. HEK293T/17 (cat# CRL-11268), I1 mouse hybridoma (cat# CRL-2700) and Vero E6 cells (cat# CRL-1586) were from ATCC.

FreeStyle 293-F cells and were cultured in serum-free FreeStyle 293 Expression Medium (GIBCO, cat# 12338026) at 37 °C, 10% CO_2_, 115 rpm. Expi293F cells were cultured in Expi293 Expression Medium (GIBCO, cat# A14635) at 37 °C, 8% CO_2_, 125 rpm. HEK293T/17 cells and Vero E6 cells were cultured in 10% Fetal Bovine Serum (FBS, GIBCO cat# 16140071) supplemented Dulbecco’s Modified Eagle Medium (DMEM, ATCC cat# 30-2002) at 37 °C, 5% CO_2_. Cell lines were not specifically authenticated.

### METHOD DETAILS

#### Protein samples expression and purification

SARS-CoV-2 S2P spike was produced as described in (Wrapp et al., 2020b). Protein expression was carried out in Human Embryonic Kidney (HEK) 293 Freestyle cells (Invitrogen) in suspension culture using serum-free media (Invitrogen) by transient transfection using polyethyleneimine (Polysciences). Cell growths were harvested four days after transfection, and the secreted protein was purified from supernatant by nickel affinity chromatography using Ni-NTA IMAC Sepharose 6 Fast Flow resin (GE Healthcare) followed by size exclusion chromatography on a Superdex 200 column (GE Healthcare) in 10 mM Tris, 150 mM NaCl, pH 7.4.

Biotinylated N-terminal domain of SARS-CoV-2 spike (NTD, residues 1-330) was cloned into the pLEXm mammalian cell expression vector proceeded by a BiP signal peptide and in frame with a C-terminal 6X-His tag and an Avi-tag (GLNDIFEAQKIEWHE). The NTD-Avi tag-expression plasmid was transiently co-transfected with the pVRC8400 plasmid encoding the biotin-Ligase BirA from *E. coli* (Lys2-Lys321) into HEK293 cells suspension culture in serum-free media using polyethyleneimine transfectant. The NTD-expression plasmid and BirA plasmid were mixed at a 10:1 ratio for transfection and 3 hrs post-transfection the media was supplemented with 50 μM Biotin (Sigma). Media was harvested 4 days after transfection and the secreted protein purified using Ni-NTA IMAC Sepharose 6 Fast Flow resin (Cytiva) followed by size exclusion chromatography (SEC) on Superdex 200 (Cytiva) in 10 mM, Tris pH 7.4, 150 mM NaCl.

NTD-directed monoclonal antibodies 5-7, 4-8, and 5-24 for neutralization assays were expressed and purified as described in Liu et al., 2020a. For SPR experiments, Fabs fragments were produced by digestion of IgGs with immobilized papain at 37 °C for 4 h in 50 mM phosphate buffer, 120 mM NaCl, 30 mM cysteine, 1 mM EDTA, pH 7. The resulting Fabs were either purified from Fc by affinity chromatography on protein A (5-7, 4-8) or used as Fab/Fc mixture (5-24).

Monoclonal antibody 5-7 for cryo-EM experiments was expressed and purified as Fab: VHCH1 with a C-terminal His-tag (His_8_) and LC were constructed separately into the gWiz expression vector, and then co-transfected and expressed in Expi293. Five days after transfection, supernatants were harvested and 5-7 Fab was purified by nickel affinity chromatography using Ni-NTA agarose (Invitrogen cat# R901-15). Fab purity was assessed by SDS-PAGE; all Fabs were buffer-exchanged into 10 mM Tris, 150 mM, pH 7.4.

#### Cryo-EM sample preparation

The final sample for EM analysis of the 5-7 in complex with SARS-CoV-2 S2P spike was produced by mixing the Fab and spike in a 1:9 molar ratio, with a final trimer concentration of 0.33 mg/mL, followed by incubation on ice for 1 hr. The final buffer was 10 mM sodium acetate, 150 mM NaCl, 0.005% (w/v) n-Dodecyl β-D-maltoside, pH 4.5. Cryo-EM grids were prepared by applying 2 μL of sample to a freshly glow-discharged carbon-coated copper grid (CF 1.2/1.3 300 mesh); the sample was vitrified in liquid ethane using a Vitrobot Mark IV with a wait time of 30 s and a blot time of 3 s.

#### Cryo-EM data collection, processing and structure refinement

Cryo-EM data were collected using the Leginon software (Suloway et al., 2005) installed on a Titan Krios electron microscope operating at 300 kV, equipped with a Gatan K3-BioQuantum direct detection device. The total dose was fractionated for 3 s over 60 raw frames. Motion correction, CTF estimation, particle extraction, 2D classification, ab initio model generation, 3D refinements and local resolution estimation for all datasets were carried out in cryoSPARC 3.2(Punjani et al., 2017); particles were picked using Topaz (Bepler et al., 2019). Bayesian polishing in RELION was performed on the final set of particles (Scheres, 2012; Zivanov et al., 2019). The final 3D reconstruction was obtained using non-uniform refinement with C1 symmetry in cryoSPARC.

SARS CoV-2 S2P spike density was modeled using PDB entry 7L2E (Cerutti et al., 2021a), as initial template. The initial model for 5-7 Fab variable region was obtained using the SAbPred server (Dunbar et al., 2016). Automated and manual model building were iteratively performed using real space refinement in Phenix (Adams et al., 2004) and Coot (Emsley and Cowtan, 2004) respectively. Half maps were provided to Resolve Cryo-EM tool in Phenix to support manual model building. Geometry validation and structure quality assessment were performed using EMRinger (Barad et al., 2015) and Molprobity (Davis et al., 2004). Map-fitting cross correlation (Fit-in-Map tool) and figures preparation were carried out using PyMOL, UCSF Chimera (Pettersen et al., 2004) and Chimera X (Pettersen et al., 2021). A summary of the cryo-EM data collection, reconstruction and refinement statistics is shown in Table S1.

#### Authentic SARS-CoV-2 microplate neutralization

The SARS-CoV-2 viruses USA-WA1/2020 (WA1), USA/CA_CDC_5574/2020 (B1.1.7), hCoV-19/South Africa/KRISP-EC-K005321/2020 (B1.351), hCoV-19/Japan/TY7-503/2021 (P.1) and hCoV-19/USA/CA (B.1.429) were obtained from BEI Resources (NIAID, NIH) and propagated for one passage using Vero E6 cells. hCoV-19/USA/NY-NP-DOH1/2021 was isolated and sequence was verified (Annavajhala et al., 2021). Virus infectious titer was determined by an end-point dilution and cytopathic effect (CPE) assay on Vero E6 cells as described previously(Liu et al., 2020a).

An end-point dilution microplate neutralization assay was performed to measure the neutralization activity of antibodies. In brief, antibodies were subjected to successive 5-fold dilutions starting from 50 μg/mL. Triplicates of each dilution were incubated with SARS-CoV-2 at a MOI of 0.1 in EMEM with 7.5% inactivated fetal calf serum (FCS) for 1 hour at 37 °C. Post incubation, the virus-antibody mixture was transferred onto a monolayer of Vero E6 cells grown overnight. The cells were incubated with the mixture for ~70 hours. CPE of viral infection was visually scored for each well in a blinded fashion by two independent observers. The results were then reported as percentage of neutralization at a given antibody dilution.

#### Pseudovirus neutralization assays

Plasmids encoding the single and combination mutations found in variants were generated by Quikchange II XL site-directed mutagenesis kit (Agilent). Recombinant Indiana VSV (rVSV) expressing different SARS-CoV-2 spike variants were generated as previously described (Liu et al., 2020a; Wang et al., 2021c). Briefly, HEK293T cells were grown to 80% confluency before transfection with the spike gene using Lipofectamine 3000 (Invitrogen). Cells were cultured overnight at 37 °C with 5% CO_2_, and VSV-G pseudo-typed ΔG-luciferase (G*ΔG-luciferase, Kerafast) was used to infect the cells in DMEM at a MOI of 3 for 2 hours before washing the cells with 1X DPBS three times. The next day, the transfection supernatant was harvested and clarified by centrifugation at 300 g for 10 min. Each viral stock was then incubated with 20% I1 hybridoma (anti-VSV-G, ATCC: CRL-2700) supernatant for 1 hour at 37 °C to neutralize contaminating VSV-G pseudo-typed ΔG-luciferase virus before measuring titers and making aliquots to be stored at −80 °C.

Neutralization assays were performed by incubating pseudoviruses with serial dilutions of mAbs or heat-inactivated plasma or sera, and scored by the reduction in luciferase gene expression as previously described (Wang et al., 2021c). Briefly, Vero E6 cells (ATCC) were seeded in 96-well plates (2 ×104 cells per well). Pseudoviruses were incubated with serial dilutions of the test samples in triplicate for 30 min at 37 °C. The mixture was added to cultured cells and incubated for an additional 16 hrs. Luminescence was measured using Luciferase Assay System (Promega), and IC50 was defined as the dilution at which the relative light units were reduced by 50% compared with the virus control wells (virus + cells) after subtraction of the background in the control groups with cells only. The IC50 values were calculated using a five-parameter dose-response curve in GraphPad Prism.

#### SPR experiments

SPR binding assays for Fabs 5-7, 4-8 and 5-24 binding to NTD were performed using a Biacore T200 biosensor, equipped with a Series S SA chip, at 25 °C. NTD carrying a C-terminal Avi-tag, was captured over a single streptavidin flow cell at approximately 700 RU. A streptavidin surface was used as a reference flow cell to remove bulk shift changes from the binding signal.

Binding experiments for each of the three Fabs binding to NTD were performed in a ligand-free buffer that consisted of HBS-T pH 7.4 (10 mM HEPES pH 7.4, 150 mM NaCl, 0.05% (v/v) Tween-20), and in buffers each supplemented with ligands that have been reported to bind to the NTD hydrophobic pocket such as 1) 0.2 μM biliverdin, 0.5% (v/v) DMSO, 2) 10 μM bilirubin, 1% (v/v) DMSO and 3) 0.01% (v/v) Polysorbate-80 (P-80). Fabs 5-7 and 5-24 were tested using a three-fold dilution series ranging from 1.1-270 nM, and 4-8 was analyzed at concentrations ranging 1.1-90 nM, using also a three-fold dilution series. Each Fab was tested in order of increasing protein concentration, in duplicate. The association and dissociation rates were each monitored for 120 s and 600 s respectively, at 50 μL/min. The NTD surface was regenerated using two-consecutive 10 s pulse of 15 mM H3PO4 at a flow rate of 100 μL/min, followed by a 60 s buffer wash at the same flow rate. Blank buffer cycles were performed by injecting running buffer instead of Fab to remove systematic noise from the binding signal. The data was processed and fit to 1:1 interaction model using the Scrubber 2.0 (BioLogic Software).

#### Antibody gene assignments and genetic analyses

The 13 NTD-directed SARS-COV-2 neutralizing and 1 non-neutralizing antibodies were collected from six publications. We annotated these antibodies using IgBLAST-1.16.0 with default parameters (Ye et al., 2013). For antibody 5-7, the N-addition, D gene, and P-addition regions were annotated by IMGT V-QUEST (Brochet et al., 2008). The gene specific substitution profile of IGHV1-46 and IGKV1-9 were download from cAb-Rep database (Guo et al., 2019).

#### Sequence entropy comparison

The residues of 5-7 paratope were obtained by PDBePISA (Krissinel and Henrick, 2007), with the default parameters. Per residue sequence entropy was download from the next strain database (Hadfield et al., 2018), which enabled by data from GISAID (https://www.gisaid.org/). The p value among different paratope groups were calculated by Kolmogorov–Smirnov test in R, with the significant level set as p < 0.05. The geometric mean and 95% CI of per residue sequence entropy were calculated and plotted by GraphPad Prism version 9. The visualization of sequence entropy was displayed by PyMOL version 2.3.2.

### QUANTIFICATION AND STATISTICAL ANALYSIS

The statistical analyses for the pseudovirus neutralization assessments were performed using GraphPad Prism. Cryo-EM data were processed and analyzed using cryoSPARC and RELION. The SPR data were fitted using Biacore Evaluation Software and Scrubber. Cryo-EM and crystallographic structural statistics were analyzed using Phenix, Molprobity, EMringer and Chimera. The correlations were performed in R. Statistical details of experiments are described in Method Details or figure legends.

