## Supplemental Figures and Tables for "Neutralizing antibody 5-7 defines a distinct site of vulnerability in SARS-CoV-2 spike N-terminal domain"

A

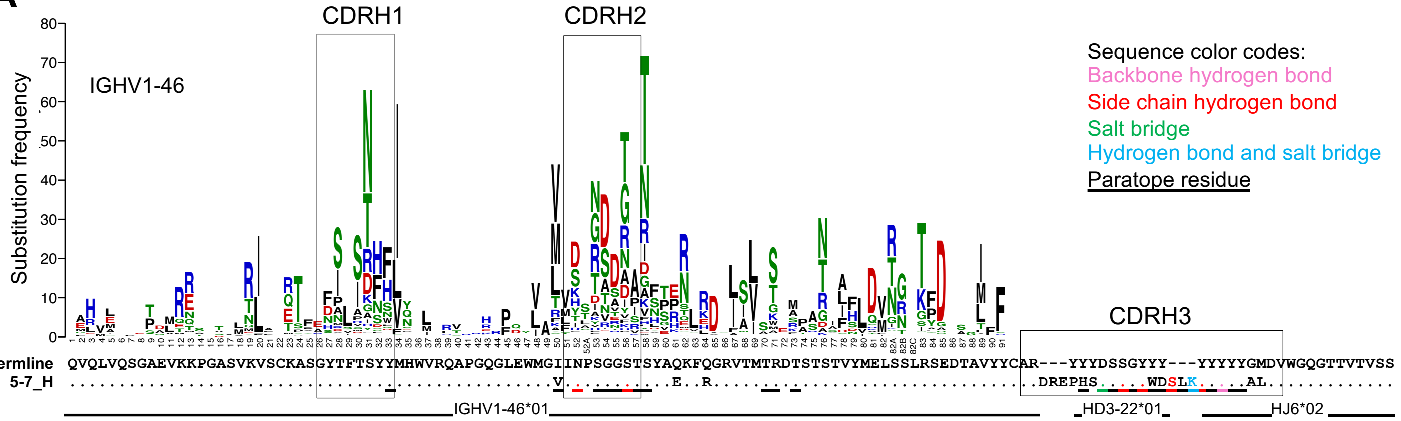


B

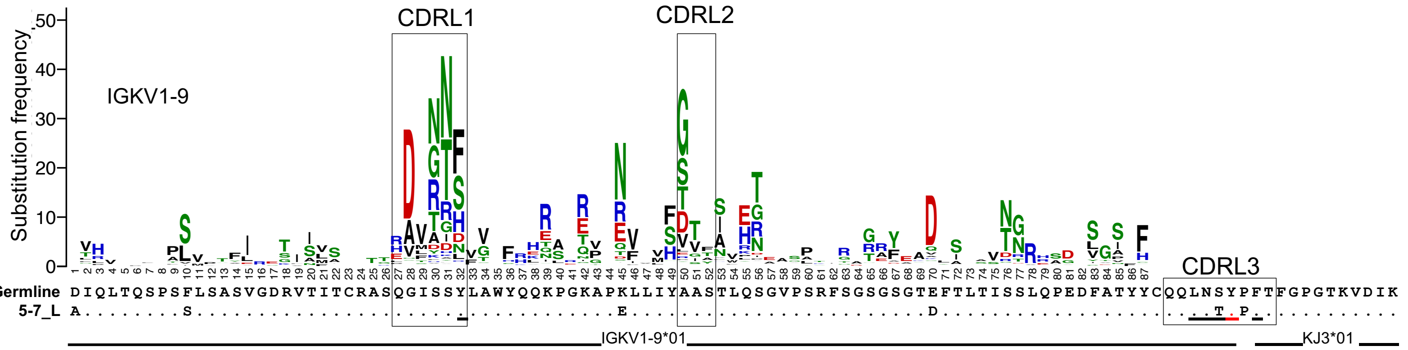


C

| mAb | HV | HJ | LV | LJ | CDRH3 | CDRL3 | Structure | Reference | Neutralizing SARS-CoV-2 | Targets NTD supersite |
|---|---|---|---|---|---|---|---|---|---|---|
| 5-7 | IGHV1-46 | IGHJ6 | IGKV1-9 | IGKJ3 | ARDREPHSDSSGYWDSLKYYYYYAL | QQLNTYPFT | 7N01 | This paper | Yes | No |
| 4A8 | IGHV1-24 | IGHJ6 | IGKV2-24 | IGKJ3 | ATSTAVAGTPDLFDYYYGMDV | TQATQFPYT | 7C2L | Chi et al., 2020 | Yes | Yes |
| FC05 | IGHV1-24 | IGHJ5 | IGLV1-51 | IGLJ3 | ATTTPFSSSYWFDP | GTWDSSLSAVV | 7CWS | Wang et al., 2020 | Yes | Yes |
| CM25 | IGHV1-24 | IGHJ5 | IGLV1-51 | ND | ATGPAVRRGSWFDP | ND | 7M8J | Voss et al., 2020 | Yes | Yes |
| 1-87 | IGHV1-24 | IGHJ6 | IGLV2-14 | IGLJ1 | ATGIAVIGPPPSTYYYYGMDV | SSYTSSSTYV | 7L2D | Cerutti et al., 2021 | Yes | Yes |
| 2-17 | IGHV1-69 | IGHJ5 | IGKV3-15 | IGKJ4 | ARGVGYRGVIPLNWFDP | QQYNNWPPFT | 7LQW | Cerutti et al., 2021 | Yes | Yes |
| 2-51 | IGHV1-24 | IGHJ4 | IGLV2-8 | IGLJ3 | ATGWAYKSTWYFGY | SSYAGSRMG | 7L2C | Cerutti et al., 2021 | Yes | Yes |
| 4-8 | IGHV1-69 | IGHJ6 | IGLV3-1 | IGLJ3 | ASLQTVDTAIEKYYGMDV | QAWDSSTAV | 7LQV | Cerutti et al., 2021 | Yes | Yes |
| 4-18 | IGHV3-30 | IGHJ5 | IGLV3-25 | IGLJ3 | AKDSGYNYGYSWFDP | QSTDNSGTYPNWV | 7L2E | Cerutti et al., 2021 | Yes | Yes |
| 5-24 | IGHV3-33 | IGHJ6 | IGKV3-20 | IGKJ4 | ARDPRDYYDFWSGYDYYYGLDV | QQYGSSGALT | 7L2F | Cerutti et al., 2021 | Yes | Yes |
| S2L28 | IGHV3-21 | IGHJ4 | IGLV2-14 | IGLJ3 | ARDGNAYKWLLAENVRFDY | SSYTSSSTPNWV | 7LXZ | McCallum et al., 2021 | Yes | Yes |
| S2X333 | IGHV3-33 | IGHJ4 | IGLV3-21 | IGLJ3 | ARAFPDSSSWSGFTIDY | QVWDSGSDQVI | 7LXY | McCallum et al., 2021 | Yes | Yes |
| S2M28 | IGHV3-33 | IGHJ4 | IGLV3-25 | IGLJ3 | ARAVAGEWYFDY | QSADSIGSSWV | 7LY2 | McCallum et al., 2021 | Yes | Yes |
| DH1050.1 | IGHV1-24 | IGHJ5 | IGLV2-8 | IGLJ1 | ATGSPFGVVTDWFDP | SSYAGSNNPYV | 7LCN | Li et al., 2021 | Yes | Yes |
| DH1052 | IGHV1-69-2 | IGHJ4 | IGKV3-20 | IGKJ1 | ATSSGPSRLCGGGSCYHSFDY | QQYGSSPTWT | 7LAB | Li et al., 2021 | No | No |

**Figure S1. Sequence alignment for 5-7 with their corresponding germline genes, Related to Figure 1.**

(A) 5-7 heavy chain aligned with VH1-46*01, the gene-specific substitution profile (GSSP) showing somatic hypermutation probabilities for VH1-46 gene.

(B) 5-7 light chain aligned with IG KV1-9*01, the gene-specific substitution profile (GSSP) showing somatic hypermutation probabilities for VK1-9 gene. The sequence color code is the same as panel A.

(C) Published structures of SARS-CoV-2 antibodies targeting NTD.

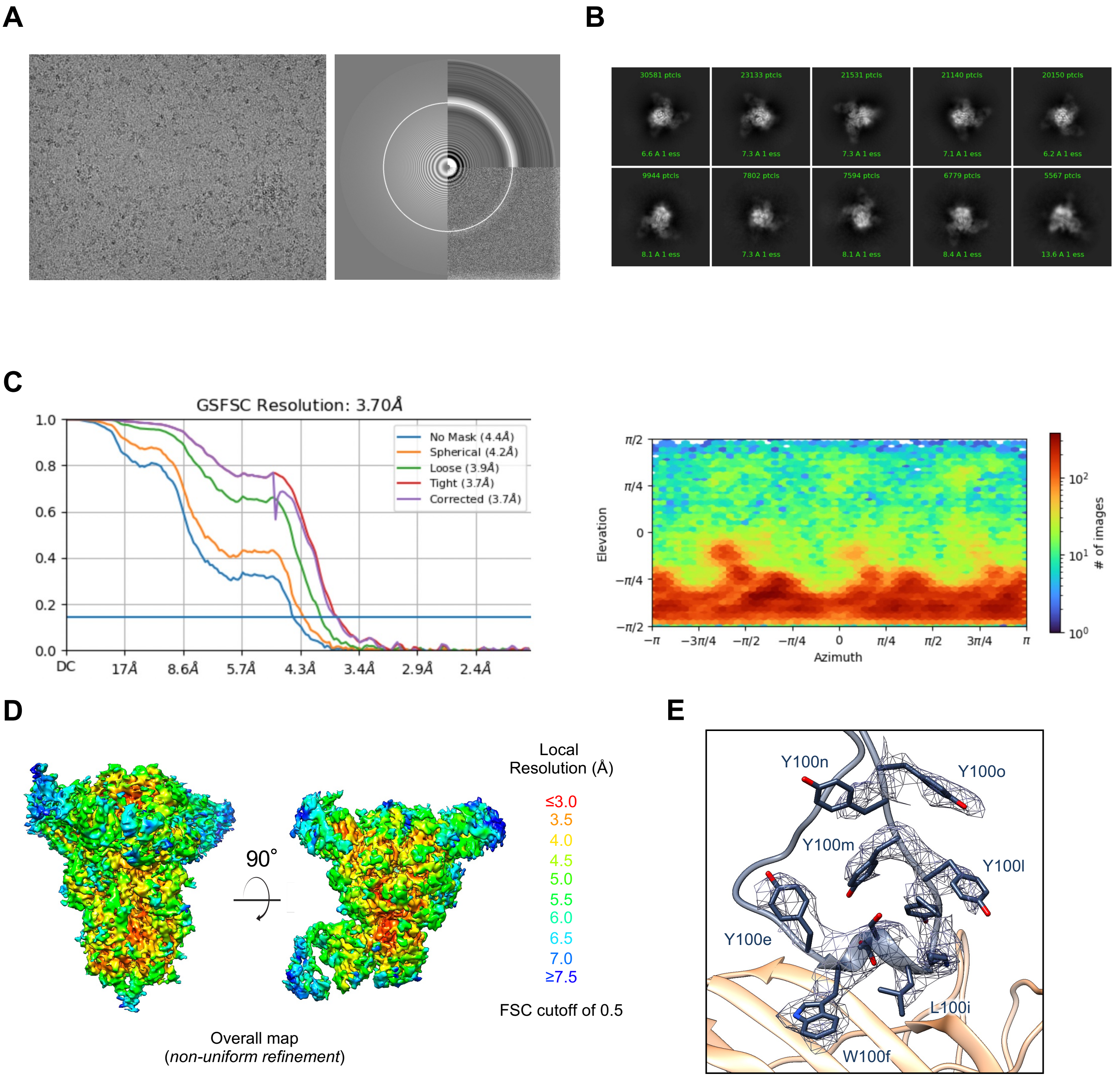


**Figure S2. Cryo-EM details of 5-7 Fab in complex with SARS-CoV-2 S2P spike, Related to Figure 1.**

(A) Representative micrograph and CTF of the micrograph are shown.

(B) Representative 2D class averages are shown.

(C) The gold-standard Fourier shell correlation resulted in a resolution of 3.70 Å for the overall map using non-uniform refinement (left panel); the orientations of all particles used in the final refinement are shown as a heatmap (right panel).

(D) The local resolution of the final overall map is shown from two orthogonal views, generated through cryoSPARC using an FSC cutoff of 0.5. The highest resolution is observed at the NTD interface with 5-7.

(E) Representative density is shown for the CDR H3 loop of 5-7 contacting NTD; the contour level is 0.45 (1.1σ). CDR H3 carbon atoms are colored in dark blue, oxygen in red, nitrogen in blue; NTD is colored in orange.

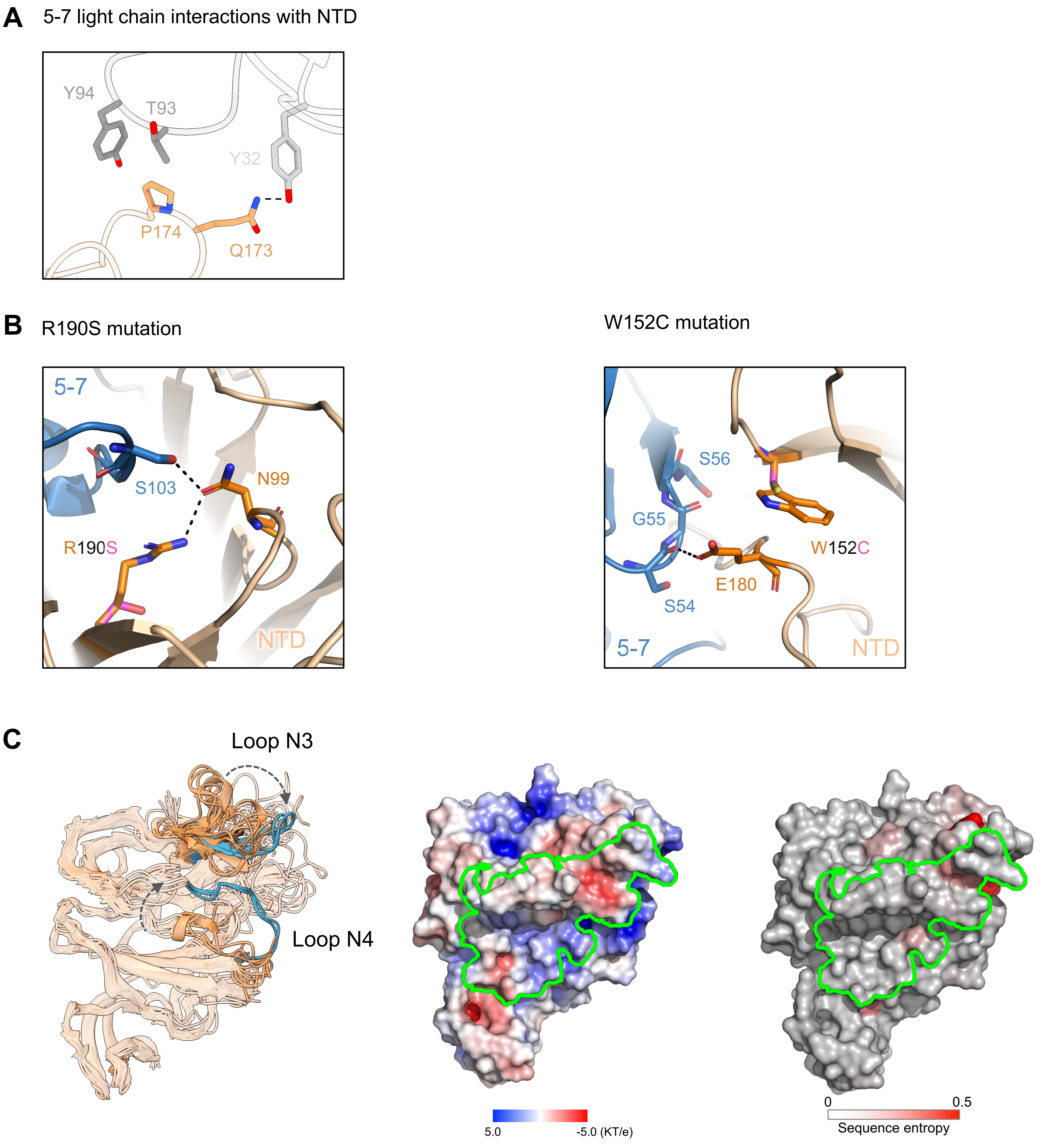


**Figure S3. Additional aspects of 5-7 binding to NTD, Related to Figures 1, 2 and 4.**

(A) Light chain contacts observed in the 5-7:spike complex. NTD is shown in orange, CDR H3 in dark gray, CDR L1 in light gray.

(B) Structural explanation for indirect effect of R190S (left panel) and W152C (right panel) on 5-7 neutralization. The hydrogen bonds are represented as black dots.

(C) Left panel: the N4 loop conformation adopts a "closed" conformation in most NTD-directed antibody complexes, while the "open" conformation is observed only for 5-7, S2L28 and ligand-free at pH 8, with their N3 and N4 loops colored in blue. The N4 loop conformation is coupled to the positioning of N3 loop (gray arrows). Middle panel: electrostatic potential and epitope outline of 5-7 on NTD. Right panel: sequence conservation of NTD.

**A**

| Fab | condition | $k_a$ ($M^{-1}s^{-1}$) | $k_d$ ($s^{-1}$) | $K_D$ (nM) |
|---|---|---|---|---|
| 5-7 | ligand-free | $1.170(9)x10^4$ | $2.49(1)x10^{-4}$ | 6.94(9) |
| | 0.2 μM biliverdin | $2.33(5)x10^4$ | $8.6(1)x10^{-4}$ | 37.1(4) |
| | 10 μM bilirubin | $2.56(4)x10^4$ | $5.80(4)x10^{-4}$ | 22.6(2) |
| | 0.01% (v/v) P-80 | $3.46(3)x10^4$ | $4.56(2)x10^{-4}$ | 13.18(7) |
| 4-8 | ligand-free | $1.479(3)x10^5$ | $3.774(7)x10^{-3}$ | 25.51(2) |
| | 0.2 μM biliverdin | $1.664(3)x10^5$ | $4.035(5)x10^{-3}$ | 24.24(2) |
| | 10 μM bilirubin | $1.630(3)x10^5$ | $3.914(6)x10^{-3}$ | 24.02(2) |
| | 0.01% (v/v) P-80 | $1.571(3)x10^5$ | $3.664(5)x10^{-3}$ | 23.32(2) |
| 5-24 | ligand-free | $6.85(1)x10^4$ | $2.173(3)x10^{-4}$ | 3.170(6) |
| | 0.2 μM biliverdin | $7.08(1)x10^4$ | $3.192(4)x10^{-4}$ | 4.511(7) |
| | 10 μM bilirubin | $6.48(1)x10^4$ | $2.951(3)x10^{-4}$ | 4.553(6) |
| | 0.01% (v/v) P-80 | $6.71(2)x10^4$ | $2.854(5)x10^{-4}$ | 4.25(1) |

**B**

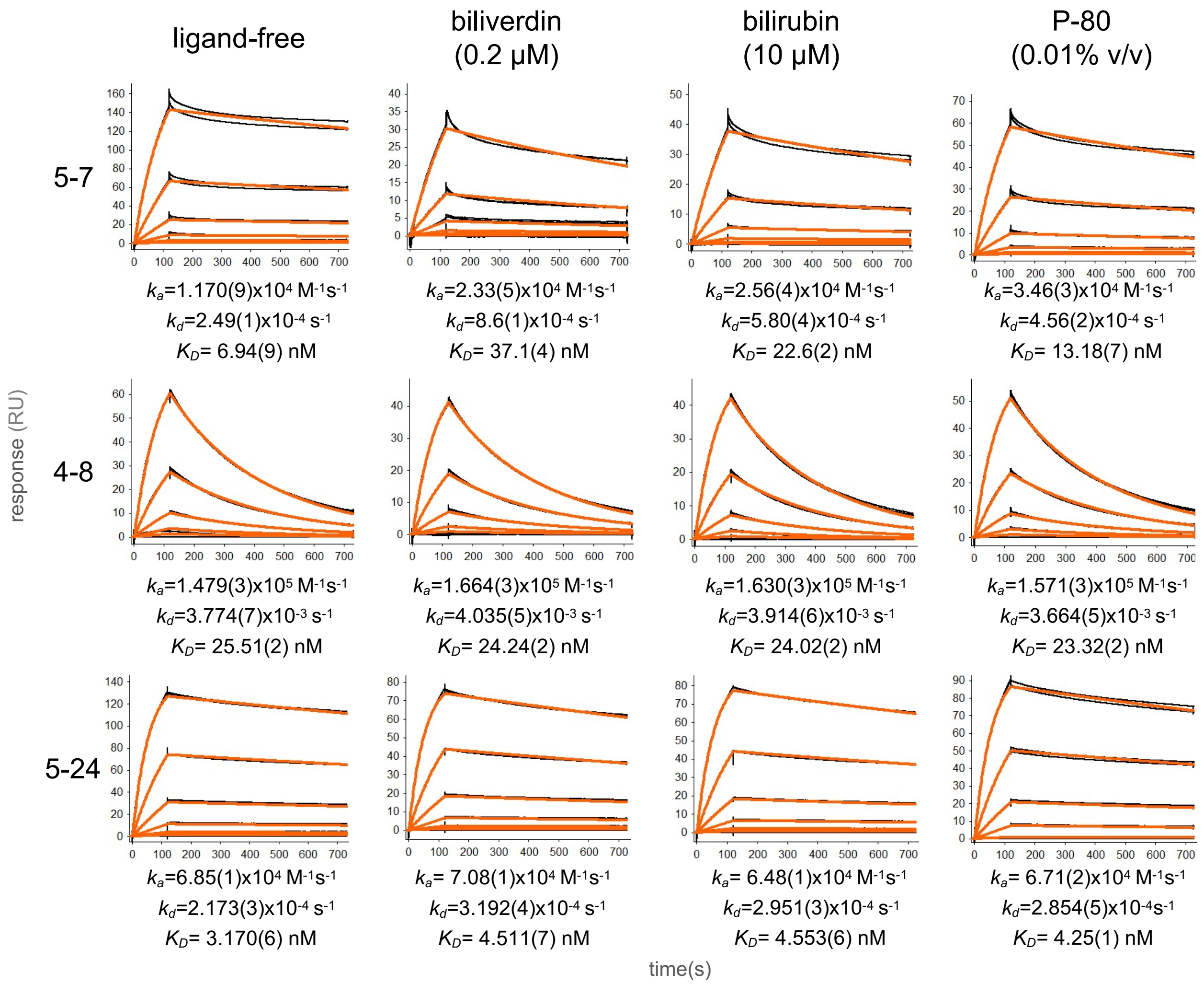


**Figure S4. SPR results showing Fabs binding to NTD in the presence of ligands reported to bind to the NTD hydrophobic pocket**

(A) Table of kinetics parameters for Fabs 5-7, 4-8 and 5-24 binding to NTD, observed without ligand and in the presence of biliverdin, bilirubin and polysorbate-80 (P-80). The number in brackets indicates the error of the fit at the last significant figure.

(B) SPR experimental data showing the binding profiles for Fabs 5-7, 4-8 and 5-24 in the buffer conditions described in a. The black traces represent the experimental data, and the red lines represent the fit to a 1:1 interaction model.

**Table S1. Cryo-EM Data Collection and Refinement Statistics, Related to Figure 1.**

| SARS-CoV-2 S2P complex | 5-7 Fab |
|---|---|
| **EMDB ID** | EMD-24097 |
| **PDB ID** | 7N01 |
| Data Collection | |
| Microscope | FEI Titan Krios |
| Voltage (kV) | 300 |
| Electron dose ($e^-/Å^2$) | 41.92 |
| Detector | Gatan K3 BioQuantum |
| Pixel Size (Å) | 1.07 |
| Defocus Range (μm) | -0.8/-2.5 |
| Magnification | 81000 |
| Reconstruction | |
| Software | cryoSPARC v3.2.0 |
| Particles | 150,834 |
| Symmetry | C1 |
| Box size (pix) | 400 |
| Resolution (Å) ($FSC_{0.143}$) | 3.70 |
| Refinement | |
| Software | Phenix 1.19 |
| Protein residues | 3416 |
| Chimera CC | 0.79 |
| EMRinger Score | 1.2 |
| R.m.s. deviations | |
| Bond lengths (Å) | 0.006 |
| Bond angles (°) | 1.33 |
| Validation | |
| Molprobity score | 1.58 |
| Clash score | 4.77 |
| Favored rotamers (%) | 99.8 |
| Ramachandran | |
| Favored regions (%) | 95.2 |
| Allowed regions (%) | 4.8 |
| Disallowed regions (%) | 0 |
